# Perturbation biology models predict c-Myc as an effective co-target in RAF inhibitor resistant melanoma cells

**DOI:** 10.1101/008201

**Authors:** Anil Korkut, Weiqing Wang, Emek Demir, Bülent Arman Aksoy, Xiaohong Jing, Evan Molinelli, Özgün Babur, Debra Bemis, David B. Solit, Christine Pratilas, Chris Sander

## Abstract

Systematic prediction of cellular response to perturbations is a central challenge in biology, both for mechanistic explanations and for the design of effective therapeutic interventions. We addressed this challenge using a computational/experimental method, termed perturbation biology, which combines high-throughput (phospho)proteomic and phenotypic response profiles to targeted perturbations, prior information from signaling databases and network inference algorithms from statistical physics. The resulting network models are computationally executed to predict the effects of tens of thousands of untested perturbations. We report cell type-specific network models of signaling in RAF-inhibitor resistant melanoma cells based on data from 89 combinatorial perturbation conditions and 143 readouts per condition. Quantitative simulations predicted c-Myc as an effective co-target with BRAF or MEK. Experiments showed that co-targeting c-Myc, using the BET bromodomain inhibitor JQ1, and the RAF/MEK pathway, using kinase inhibitors is both effective and synergistic in this context. We propose these combinations as pre-clinical candidates to prevent or overcome RAF inhibitor resistance in melanoma.

## Introduction

### Drug resistance in cancer treatment

The Inhibition of key oncogenes with target specific agents elicits dramatic initial response in some cancers such as prostate cancer and melanoma (Bollag et al., 2010; Clegg et al., 2012). Even for the most successful single agent targeted therapies, however, drug resistance eventually emerges leading to rapid progression of metastatic disease (Garraway and Janne, 2012). The mechanism of drug resistance may involve selection of resistant sub-clones, emergence of additional genomic alterations, and compensating interactions between alternative signaling pathways (Choi et al., 2007; Huang et al., 2011; Wagle et al., 2013). One potential solution to overcome drug resistance is to combine targeted drugs to block potential escape routes (Fitzgerald et al., 2006). Therefore, there is currently a need for systematic strategies to develop effective drug combinations.

### Drug resistance in melanoma

Targeted therapy has been particularly successful in treatment of melanoma. *BRAFV600E* gain-of-function mutation is observed in ∼50% of melanomas (Davies et al., 2002). Direct inhibition of BRAFV600E by the RAF inhibitor (RAFi) vemurafenib and inhibition of the downstream MEK and ERK kinases have yielded response rates of more than 50% in melanoma patients with this mutation (Chapman et al., 2011; Flaherty et al., 2012). At the cellular level, inhibition of the ERK pathway leads to changes in expression of a set of critical cell cycle genes (e.g., *CCND1, MYC, FOS*) and feedback inhibitors of ERK signaling (e.g., *DUSP, SPRY2*) (Pratilas et al., 2009). Resistance to vemurafenib emerges after a period of ∼7 months in tumors that initially responded to single-agent therapy (Chapman et al., 2011; Sosman et al., 2012). Multiple RAFi and MEKi (e.g., PD-0325901, Trametinib) resistance mechanisms, which may involve alterations in NRAS/ERK pathway (e.g., NRAS mutations, switching between RAF isoforms) or parallel pathways (e.g., PTEN loss), have been discovered in melanoma (Johannessen et al., 2010; Nazarian et al., 2010; Poulikakos et al., 2010; Xing et al., 2012).

The alterations associated with drug resistance may pre-exist alone, in combinations or emerge sequentially and vary substantially between patients (Van Allen et al., 2014). Effective drug combinations may target diverse resistance mechanisms. Despite anecdotal success, conventional methods and combinatorial drug screens generally fail to come up with effective combinations due to the genomic complexity and heterogeneity of tumors (Zhao et al., 2014). In order to more effectively nominate drug combinations, we propose to employ system-wide models that cover interactions between tens to hundreds of signaling entities and can describe and predict cellular response to multiple interventions. There have been prior attempts to construct such signaling models. De novo and data-driven quantitative models were able to cover only a few signaling interactions and therefore had limited predictive power (Nelander et al., 2008; Bender et al., 2011; Klinger et. al., 2014; Oates, et al. 2014).

Qualitative or discrete models can cover more interactions but typically lack the capability of generating quantitative predictions (Saez-Rodriguez et al., 2009; Breitkreutz et al., 2010; Saez-Rodriguez et al., 2011). Detailed physicochemical models derived using generic biochemical kinetics data can be quite comprehensive and quantitative but typically lack sufficient cell-type specificity required for translationally useful predictions (Chen et al., 2009).

We construct comprehensive, cell-type specific signaling models that quantitatively link drug perturbations, (phospho)proteomic changes and phenotypic outcomes (Figure 1). The models capture diverse signaling events and predict cellular response to previously untested combinatorial interventions. In order to generate the training data for network modeling, we first perform systematic perturbation experiments in cancer cells with targeted agents. Next, we profile proteomic and phenotypic response of cells to the perturbations. The cell type-specific response data serve as the input for network inference. We also incorporate prior pathway information from signaling databases to narrow the parameter search space and improve the accuracy of the models. For this purpose, we developed a computational tool [Pathway Extraction and Reduction Algorithm (PERA)] that automatically extracts priors from the Pathway Commons signaling information resource (Cerami et al., 2011; Demir et al., 2010).

**Figure 1.**
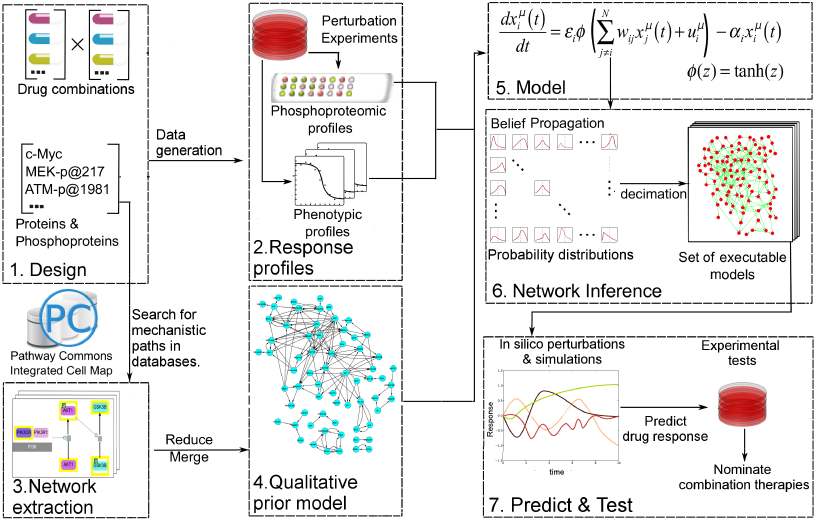
Quantitative and predictive signaling models are generated from experimental response profiles to perturbations. Perturbation biology involves systematic perturbations of cells with combinations of targeted compounds (Box 1-2), high-throughput measurements of response profiles (Box 2), automated extraction of prior signaling information from databases (Box 3-4), construction of ODE-based signaling pathway models (Box 5) with the belief propagation (BP) based network inference algorithm (Box 6) and prediction of system response to novel perturbations with the models and simulations (Box 7). The “prior extraction and reduction algorithm” (PERA) generates a qualitative prior model, which is a network of known interactions between the proteins of interest (i.e., profiled (phospho)proteins). This is achieved through a search in the Pathway Commons information resource, which integrates biological pathway information from multiple public databases (Box 3-4). In the quantitative network models, the nodes represent measured levels of (phospho)proteins or cellular phenotypes and the edges represent the influence of the upstream nodes on the time derivative of their downstream effectors. This definition corresponds to a simple yet efficient ODE-based mathematical description of models (Box 5). Our BP-based modeling approach combines information from the perturbation data (phosphoproteomic and phenotypic) with prior information to generate network models of signaling (Box 6). We execute the resulting ODE based models to predict system response to untested perturbation conditions (Box 7).

Even in the presence of large training data and priors, network inference is a difficult problem due to the combinatorial complexity (i.e., exponential expansion of the parameter search space with linear increase of parameters). For example, to infer a network model with 100 nodes using Monte Carlo based methods, we in principle would need to cover a search space that includes ∼2^(100 × 100)^ network models—a computationally impossible task. To circumvent this problem, we have adapted a network modeling algorithm based on belief propagation (BP), which replaces exhaustive one-by-one searches by a search over probability distributions (Miller et al., 2013; Molinelli, Korkut, Wang et al., 2013). The algorithm enables us to construct models that can predict response of hundreds of signaling entities to any perturbations in the space of modeled components. We use prior information (signaling interactions from databases) as soft restraints on search space, i.e., the algorithm rejects interactions that do not conform to the experimental training data. To quantitatively predict cellular response to combinatorial perturbations, we simulate the fully parameterized network models with in silico perturbations until the system reaches steady state (Figure 1 and 5). The steady state readout for each proteomic and phenotypic entity (i.e., system variables) is the predicted response to the perturbations.

We constructed cell type specific network models of signaling from perturbation experiments in RAFi-resistant melanoma cells (SkMel-133 cell line). The melanoma cells used for network modeling have a *BRAFV600E* mutation and homozygous deletions in *PTEN* and *CDKN2A*. The models quantitatively link 82 (phospho)proteomic nodes (i.e., molecular concentrations) and 12 protein activity nodes with 5 cellular phenotype nodes (e.g., cell viability). As shown by cross validation calculations, use of prior information significantly improved the predictive power of the models. Once the predictive power was established, we expanded the extent of the drug response information from a few thousand experimental data points to millions of predicted node values. Based on the predictions, which cannot be trivially deduced from experimental data, we nominated co-targeting of c-Myc and BRAF or MEK as a potential strategy to overcome RAFi-resistance. To test the prediction, we experimentally showed that the BET bromodomain inhibitor, JQ1, reduces c-Myc expression and, in synergy with RAF/MEK signaling inhibition, substantially reduces the growth of RAFi-resistant SkMel-133 cells and in this context overcomes drug resistance.

## Results

### 1. Experimental response maps from proteomic/phenotypic profiling

#### Drug Perturbation experiments in melanoma cells

We performed systematic perturbation experiments in malignant melanoma cells (Figure 2A) to generate a rich training set for network inference. The RAFi-resistant melanoma cell line SkMel-133 was treated with combinations of 12 targeted drugs (Figure 2A, Table S1 for drug targets and doses). The perturbations consisted of systematic paired combinations of individual agents and multiple doses of single agents. This procedure generated 89 unique perturbation conditions, which targeted specific pathways including those important for melanoma tumorigenesis such as ERK and PI3K/AKT (Haluska et al., 2006).

**Figure 2.**
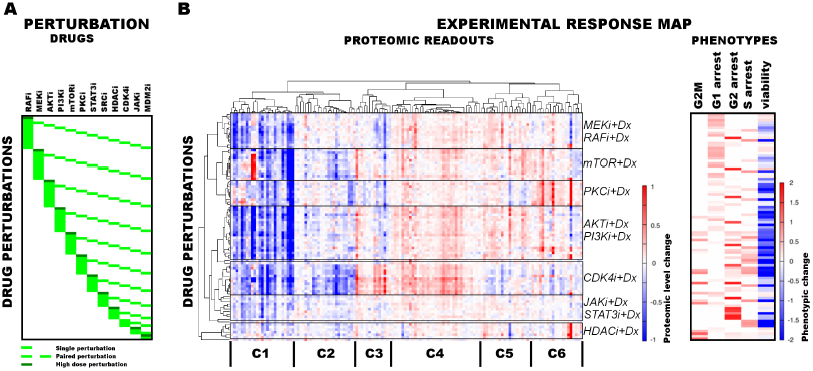
Response of melanoma cells to systematic perturbations with targeted agents. **A.** The combinatorial perturbation matrix. The melanoma cells are perturbed with combinations of targeted drugs (See Table S2 for perturbation conditions). **B.** The concentration changes in 138 proteomic entities (50 phospho, 88 total protein measurements) (left) and the phenotypic changes (right) in response to drug combinations with respect to the untreated conditions form an experimental “response map” of the cellular system. The response map reflects the functional relations between signaling proteins and cellular processes. The two-way clustering analysis of the proteomic readouts reveals distinct proteomic response signatures for each targeted drug (See Figure S2). Cell cycle progression and viability response are measured using flow cytometry and resazurin assays respectively. The cell cycle progression phenotype is quantified based on the percentage of the cells in a cell cycle state in perturbed condition with respect to the unperturbed condition. For the phenotypic readouts, the order of the perturbation conditions is same as in (A). The response values are relative to a no drug control and given as log_2_(perturbed/unperturbed).

#### Proteomic/Phenotypic Profiles

A key aspect of the data acquisition for network inference is combining the proteomic and cellular phenotypic data so that the resulting models quantitatively link the proteomic changes to global cellular responses. Toward this objective, we profiled the melanoma cells for their proteomic and phenotypic response under 89 perturbation conditions (Figure 2B-C). We used reverse phase protein arrays (RPPA) to collect drug response data for 138 proteomic (total and phospho-protein levels) entities in all conditions (Tibes et al., 2006). In parallel, we measured phenotypic responses, including cell viability and cell cycle progression (i.e., G1, S, G2, G2M arrest phenotypes) in all conditions (Figure 2B).

#### The response map

The high throughput phenotypic and proteomic profiles form a response map of cells to systematic perturbations (Figure 2). The response map provides context-specific experimental information on the associations between multiple system variables (i.e. proteomic entities) and outputs (i.e. phenotypes) under multiple conditions (i.e. perturbations). We demonstrate through hierarchical clustering of the map that each targeted drug induces a distinct proteomic response, and drugs targeting the same pathway lead to overlapping responses in the SkMel-133 cells (Figure 2B). Through a clustering-driven pathway analysis, we further show that functionally related proteins (i.e. proteins on same or related pathways) respond similarly to targeted agents (Figure 2C, Figure S2).

A description of the response map as a set of uncoupled pairwise associations between proteomic and phenotypic entities is not sufficient for achieving systematic predictions. Therefore, we built quantitative models using the experimental response map. The models describe the coupled nature of the interactions between proteins and cellular events, as well as the nonlinear dynamics of cellular responses to drug perturbations.

### 2. Quantitative and predictive network models of signaling

#### Network models

Next, we used the experimental response map (Figure 2) and the BP-based inference strategy (Figure 1) to build quantitative network models of signaling in melanoma. In the models, each node quantifies the relative response of a proteomic or phenotypic entity to perturbations with respect to the basal condition. Consequently, proteomic entities that do not respond to even a single perturbation condition, do not contribute any constraints for inference. We eliminated such entities from the network modeling with a signal-to-noise analysis (See Supplementary Methods) and included 82 of the 138 proteomic measurements in the modeling. In addition to the proteomic nodes, the models contained 5 phenotypic nodes and 12 “activity nodes”, which represent the 12 drugs and couple the effect of the targeted perturbations to the other nodes in the network. In total, network models contained 99 nodes. BP algorithm generates the probability distribution of edge strengths for every possible interaction between the nodes. The BP-guided decimation algorithm (see Methods, Figure S1) instantiates distinct network model configurations from the probability model (Montanari et al., 2007).

The mathematical formulation of the BP-based network inference is suitable for both de novo modeling (i.e. modeling with no prior information) and modeling using prior information (see Supplementary methods). Here, we used prior information to infer models with higher accuracy and predictive power compared to de novo models. Using the PERA computational tool, we derived a generic prior information model from Reactome and NCI-Nature PID databases, which were stored in Pathway Commons (Cerami et al., 2011). The prior information network contains 154 interactions spanning multiple pathways (Figure S3). Next, we added a prior prize term to the error model (Equation S1) to restrain the search space by favoring the interactions in the prior model. It is critical that the prior information does not overly restrain the inferred models and the algorithm can reject incorrect priors. To address this problem, we inferred network models using the pathway driven and randomly generated prior restraints. The statistical comparison of the networks inferred with actual (i.e., reported in databases) and random prior models indicated that the inference algorithm rejected significantly higher number of prior interactions when randomly generated priors were used for modeling (Figure S3). Finally, we integrated the experimental data and prior information to generate 4000 distinct and executable model solutions with low errors using the BP-based strategy.

### 3. Use of prior information improves the predictive power of models

#### Cross validations with and without prior information

We addressed the question whether BP-derived models have predictive power and whether use of prior information introduces further improvement. To assess the predictive power of the network models (i.e. predicting the response to untested perturbations), we performed a leave-k-out cross validation (Figure 3A). In two separate validation calculations, we withheld the response profile to every combination of either RAFi or AKTi (leave-11-out cross validation). This procedure created a partial training dataset that contains response to combinations of 11 drugs and 2 different doses of a single drug totaling to 78 unique conditions (Table S2).

**Figure 3.**
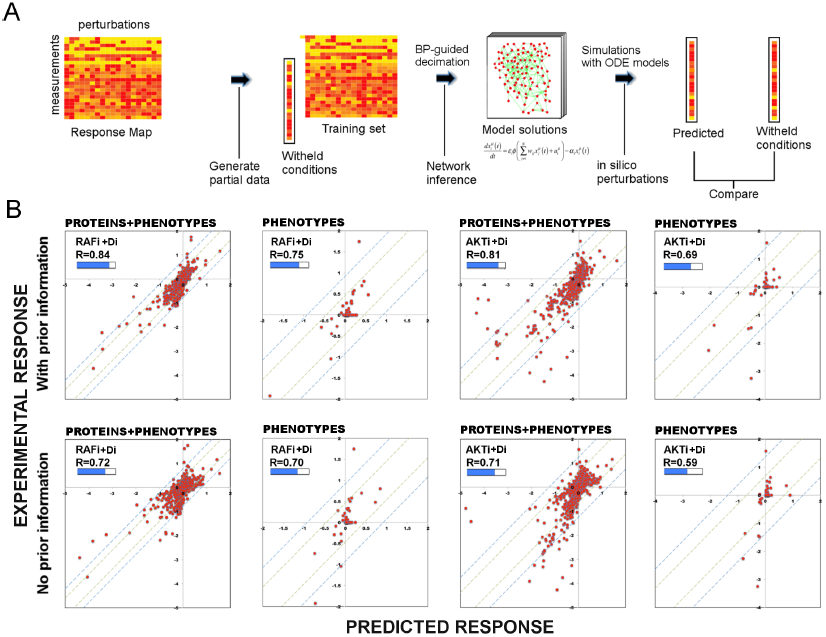
Use of prior information increases the predictive power of models. **A.** To test the predictive power of network models, a leave-11-out cross validation test is performed. Using the BP-guided decimation algorithm, 4000 network model solutions are inferred in the presence and absence of prior information using the partial response data. Resulting models are executed with in silico perturbations to predict the withheld conditions. Each experimental data point represents the read-outs from RPPA and phenotype measurements under the corresponding perturbation conditions. Each predicted data point is obtained by averaging results from simulations with in silico perturbations over 4000 model solutions. The experimental and predicted profiles are compared to demonstrate the power of network models to predict response to combinatorial drug perturbations. **B.** In all conditions, network inference with prior information leads to a higher cumulative correlation coefficient (R) and significantly improved prediction quality (RAFi p = 1 × 10^-3^, AKTi p = 5.7 × 10^-3^, unpaired t-test H_0_: ΔX^with_prior^=ΔX^w/o_prior^, ΔX=|X_exp_-X_pred_|) between experimental and predicted responses. Plots on top row: Prior information is used for network inference. Plots on bottom row: No prior information is used for network inference. Response to RAFi+{Di} (1^st^ and 2^nd^ column) and AKTi+{Di} (3^rd^ and 4^th^ column) are withheld from the training set and the withheld response is predicted. All responses (phenotypic + proteomic) (1^st^ and 3^rd^ column) and only phenotypic responses (2^nd^ and 4^th^ column) are plotted. {D_i_} denotes set of all drug perturbations combined with drug of interest.

First, we constructed both de novo (i.e. without any prior information) and prior information guided network models with each partial dataset. Next, we predicted the response by executing the models with in silico perturbations that correspond to the withheld experimental conditions. Finally, we compared the hidden and predicted response data from models generated de novo or with prior information.

#### Restraining inference with prior information improves the predictive power of models

The comparison between the predicted and the withheld experimental profiles suggests that the de novo network models have considerable predictive power and the use of prior information in modeling introduces significant improvement in the prediction quality (Figure 3B). Use of prior information increased the cumulative correlation coefficient between predicted and experimental response data from 0.72 to 0.84 and from 0.71 to 0.81 for RAFi and AKTi respectively. The demonstrated predictive power of the models suggests that the models are suitable for systematically predicting response to perturbation combinations not sampled in the training set and generating testable hypotheses that link external perturbations (e.g., targeted drugs) to cellular response.

### 4. Network models identify context dependent oncogenic signaling in melanoma

#### Network modeling and the average model

We generated quantitative network models with the complete experimental response profile and prior information to investigate oncogenic signaling in melanoma. The resulting network models resemble conventional pathway representations facilitating their comparison with the biological literature (Le Novere et al., 2009), but the interaction edges do not necessarily represent physical interactions between connected nodes. Analysis of the ensemble of network model solutions reveals that a set of strong interactions is shared by a majority of the inferred network models. On the other hand, some interactions have non-zero edge strength (Wij ≠ 0) values only in a fraction of the models (see Figure S3 for analysis of the edge distribution in models). As a first step of detailed analysis and for the purpose of intuitive interpretation, we computed an average network model (Figure 4A), which is obtained by averaging the interaction strength (Wij) for each node pair ij over all individual model solutions. This average network model highlights the interactions with high Wij values if shared by the majority of the distinct solutions. Although the average model cannot be simulated to predict system response, it is particularly useful for qualitative analysis of the inferred signaling interactions.

**Figure 4.**
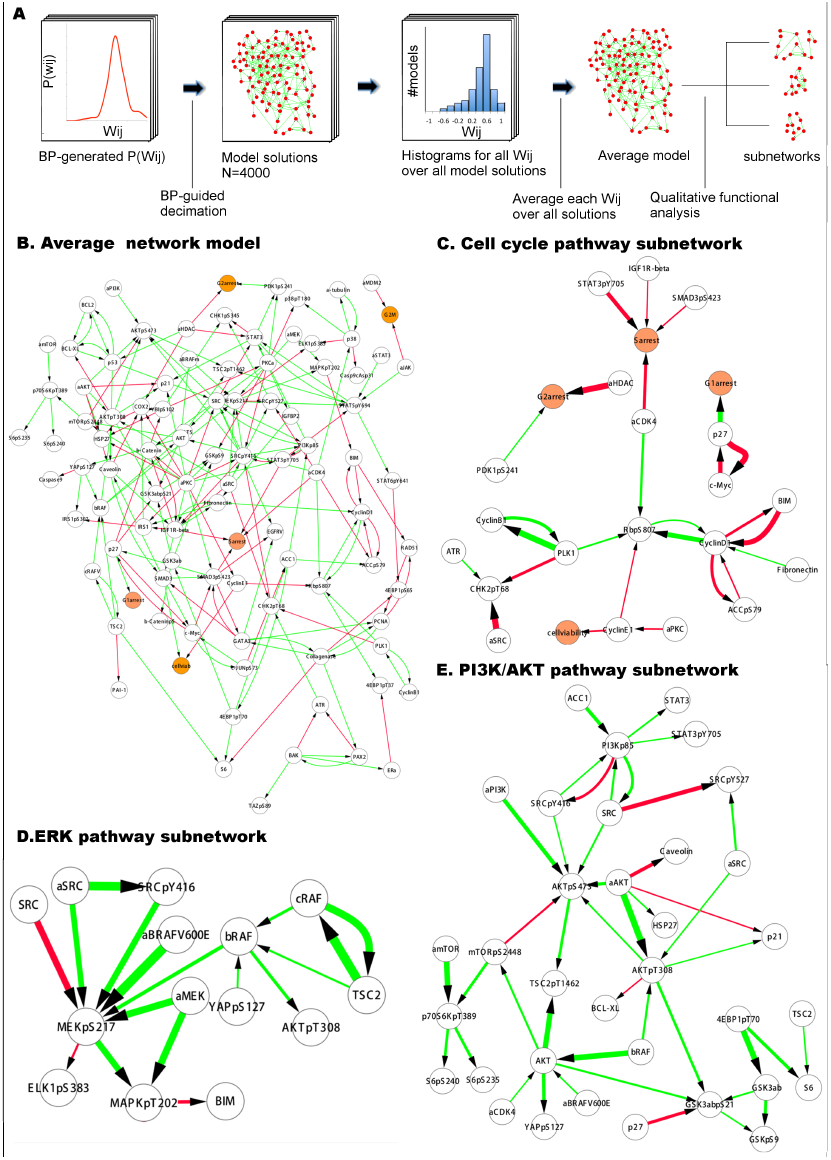
Inferred network models capture oncogenic signaling pathways in melanoma. **A.** The generation of the average model. The set of < W_ij_ > averaged over the W_ij_ in all models provide the average network model. The signaling processes are explained through qualitative analysis of the average model and its functional subnetworks (See Figure 5 for quantitative analysis). **B** The average network model contains proteomic (white) and phenotypic nodes (orange) and the average signaling interactions (> 0.2) over the model solutions. The edges between the BRAF, CRAF, TSC2 and AKTpT308 represent the cross-pathway interactions between the MAPK and PI3K/AKT pathways (See Figure S3 for analysis of edge distributions in the solution ensemble). **C.** Cell cycle signaling subnetwork contains the interactions between the cyclins, CDKs and other associated molecules (e.g., p27/Kip1). RBpS807 and cyclin D1 are the hub nodes in the subnetwork and connect multiple signaling entities. **D**. ERK subnetwork. MEKpS217 is the critical hub in this pathway and links upstream BRAF and SRC to downstream effectors such as ERK phosphorylation. **E**. In the PI3K/AKT subnetwork, the SRC nodes (i.e. phosphorylation, total level, activity) are upstream of PI3 K and AKT (total level, AKTpS473 and AKTp308) and the AKT nodes are the major hubs. Downstream of AKT, the pathway branches to mTOR, P70S6 K and S6 phosphorylation cascade and the GSK3β phosphorylation events. A negative edge originating from mTORpS2448 and acting on AKTpS473 presumably captures the well-defined negative feedback loop in the AKT pathway (O’Reilly et al., 2006). Note that nodes tagged with “a” (e.g., aBRAF) are activity nodes which couple drug perturbations to proteomic changes.

#### Global analysis of average models

The average network model provides a detailed overview of the signaling events in melanoma cells (Figure 3A). The average model contains 203 unique interactions (127 activating and 76 inhibitory interactions) between 99 signaling entities. 89 of the 154 interactions in the prior model conform to the experimental data and, therefore are accepted in the majority of the model solutions by the inference algorithm and included in the average model. Given that the average model covers interactions from multiple signaling pathways and is more complex than the pathway diagrams presented in most review papers, even the qualitative analysis of the model is highly challenging.

#### Network models capture known signaling pathways

In order to simplify the analysis of the average model solution, we fragmented the global network diagram into subnetworks (Figure 4). Each subnetwork is a simplified representation of the signaling events in canonical pathways such as those that fall into ERK, PI3K/AKT and cell cycle pathways (Figure 4C-E). The subnetwork diagrams indicate that models recapitulate many known interactions in pathways, which are important in melanoma tumorigenesis (e.g., PI3K/AKT and ERK) and nominate previously unidentified interactions (See Figure 4 legend). It is, however, not possible to predict the cellular response to untested drug perturbations through qualitative analysis of the inferred interactions. We use quantitative simulations with in silico perturbations to both decode the signaling mechanisms and more importantly systematically predict cellular response to combinatorial drug perturbations.

### 5. Combinatorial in silico perturbations generate an expanded proteomic/phenotypic response map

#### Model execution with in silico perturbations

Thanks to their ODE-based mathematical descriptions, the models can be executed to predict cellular responses to novel perturbations (Nelander et al., 2008). The systematic predictions go beyond the analysis of few particular edges in the system and capture the collective signaling mechanisms of response to drugs from the modeled pathways. We execute the parameterized model ODEs (Equation 1) with in silico perturbations acting on node (i) as a real numbered u(i) value until all the system variables (i.e. node values, {xi}) reach to steady state (Figure 5A-B).

**Figure 5.**
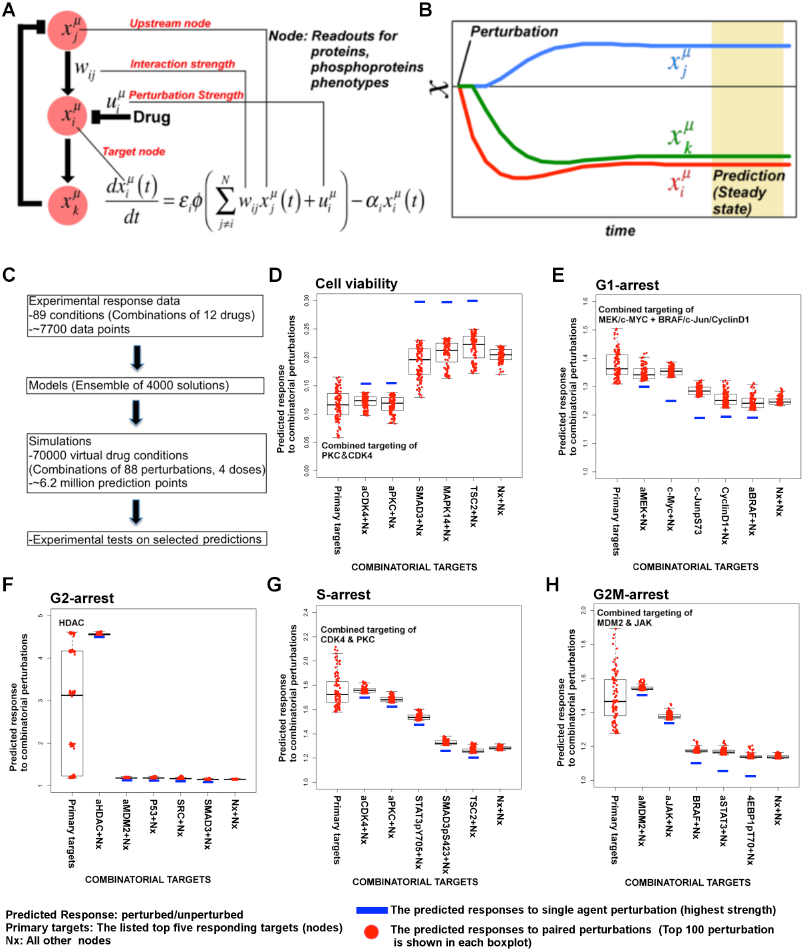
Simulations with in silico perturbations provide predictions on system response to novel perturbations. **A**. The schematic description of network simulations. The system response to paired perturbations is predicted by executing the ODE-based network models with in silico perturbations. In the ODE based models, {W_ij_} represents the set of interaction strengths and is inferred with the BP-based modeling strategy. The in silico perturbations are applied as real valued u_i_^m^ vectors. The time derivative and final concentration of any predicted node is a function of the model parameters, the perturbations and the values of all the direct and indirect upstream nodes in the models. **B**. The model equations are executed until all model variables (protein and phenotype responses) reach to steady state. The predicted response values are the averages of simulated values at steady state over 4000 distinct model solutions. **C**. The simulations expand the response map by three orders of magnitude and generate testable hypotheses. **(D-H)** The predicted phenotypic response to combinatorial in silico perturbations. Each box contains the 100 highest phenotypic responses to paired perturbations. The first box includes the response predictions for combined perturbations on primary nodes (e.g., aMEK, c-Myc for G1-arrest). The second to sixth boxes include the predicted response for combined targeting of the primary targets with all other nodes (Nx). The last box represents the predicted response data for combination of all nodes except the primaries. For the complete predicted phenotypic response see Figure S4.

#### Prediction of phenotypic responses

The simulations expand the size of the response map by three orders of magnitude from few thousand experimental response data to millions of predicted responses (Figure 5C). Once we had the predicted response profiles, we searched for specific perturbations that may induce desired phenotypic changes even when cells are treated with drugs at physiologically relevant doses. Not surprisingly, the execution of models quantitatively recaptured the experimentally observed associations between the drug perturbations and the phenotypic responses. For example, targeting PKC or CDK4 with specific kinase inhibitors leads to reduction of cell viability according to the simulations, which can also be directly observed from the experiments. However, a CDK4 and PKC inhibitors substantially reduced SkMel-133 cell viability only at high doses, as in the original perturbation experiments (> 2 µM) (Figure S5). More importantly, the simulations allowed us to identify effective perturbation combinations that cannot be trivially deduced from the experimental data (Table 1, Figure 5D-H, figures S4).

**Table 1.** Phenotypic nodes and predicted responses from simulations with in silico perturbations

| Phenotype | Immediate upstream nodes ($\langle w_{ij} \rangle$ in average model) | Predicted primary target | Predicted combination partners |
| --- | --- | --- | --- |
| **Cell viability** | SMAD3 (0.49)<br>cyclin E1 (-0.28) | aPKC<br>aCDK4 | SMAD3, 4EBP1pT70, TSC2, MAPK14/p38 |
| **G1-arrest** | p27/KIP1 (0.83)<br>PDK1pS241 (0.20) | c-Myc<br>aMEK | Cell cycle (cyclin B1, RBpS807, cyclin D1, PLK1), MAPK pathway (MAPKpT202, MEKpS217, BRAF, aBRAF), AKT, AKTpT308, RAD51, p38/MAPK14, aSRC, YB1pS102, cJUNpS73, SMAD3 |
| **G2-arrest** | aHDAC (-0.77) | aHDAC | Generic response from all nodes |
| **S-arrest** | STAT3pY705 (-0.66)<br>IGF1Rβ (-0.20) | aCDK4 | aPKC, SMAD3pS423, STAT3pS705, IGFBP2, cyclin B1, IGF1Rβ, Fibronectin |
|  | SMAD3pS423 (-0.23)<br>aCDK4 (-0.49) | aPKC | STAT3pS705 |
| **G2M** | aJAK (-0.24)<br>aMDM2 (-0.38) | aMDM2 | aJAK, aSTAT3, BRAF, 4EBPpT70 |

*\* A proteomic node corresponds to a primary target when substantial phenotypic change is predicted in response to perturbation of the node alone. The phenotypic response is further increased when the primary targets are perturbed in combination with a set of other nodes (i.e. the combination partners). Also See Figure 5D-H, Table S4 and Figure S4.*

A thorough analysis of all the predicted responses identified the top responding targets in SkMel133 cells. In particular, in silico perturbations of c-Myc lead to the strongest G1-arrest response when co-targeted with BRAF, MEK and cyclin D1 (Figure 5D-H, Figure 6A). Consistent with the vast literature on c-Myc’s role in genesis of many cancers (Dang, 2012), predictions indicated that c-Myc linked multiple pathways such as ERK and PI3K/AKT to regulation of cell cycle arrest (Table 1). As neither c-Myc nor its direct regulators were inhibited in the perturbation experiments, the predictions from the models were nontrivial and we decided to test these predictions experimentally.

**Figure 6.**
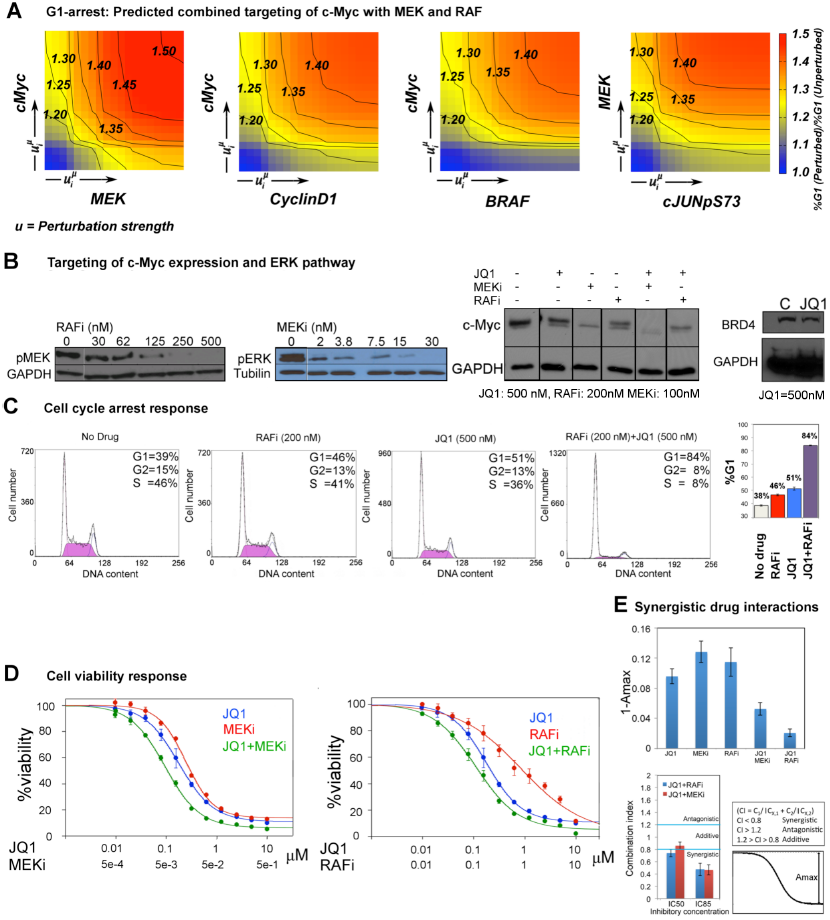
The combined targeting of c-Myc with MEK and BRAF leads to synergistic response in melanoma cells. **A**. The isobolograms of predicted G1-response to combined targeting of c-Myc with MEK, BRAF, CyclinD1 and pJUNpS73. The leftward shift of isocurves implies synergistic interactions between the applied perturbations. **u** denotes strength of in silico perturbations. **B.** RAFi inhibits MEK phosphorylation at S217 and MEKi inhibits ERK phosphorylation at T202 in a dose dependent manner (first 2 gels). Western blot showing the level of c-Myc in response to JQ1, MEKi, RAFi and their combinations (24 hours) (3^rd^ gel). c-Myc expression is targeted with JQ1 combined with MEKi or RAFi. Direct target of JQ1, BRD4 protein is expressed in both control and 500 nM JQ1 treated cells (4^th^ gel). **C.** The cell cycle progression phenotype in response to JQ1 and RAFi as measured using flow cytometry. 46% and 51% of cells are in G1 stage 24 hours after RAFi and JQ1 treatment, respectively. The combination has a synergistic effect on G1 cell cycle arrest (G1 = 84%). 39% of cells are in G1 when they are not treated with drugs. Error bars in right panel: ±SE in 3 biological replicates **D**. The drug dose-response curves of cell viability for MEKi + JQ1 (top) and RAFi + JQ1 (bottom). Cell viability is measured using the resazurin assay. Error bars: ±SEM in 3 biological replicates **E.** The synergistic interactions between JQ1 and RAFi/MEKi. 1-A_max_ is the fraction of cells alive in response to highest drug dose normalized with respect to the non-drug treated condition (top panel). Combination index (CI) quantifies the synergistic interactions between drugs (bottom left). CI is calculated at a given level of inhibition and is a measure of the fractional shift between the combination doses (C1 and C2) and the single agent’s inhibitory concentration (C_x,1_, C_x,2_).

### 6. Co-targeting c-Myc with MEK or RAF is synergistic in melanoma cells

We predicted through quantitative simulations that melanoma cells were arrested in G1-phase of the cell cycle when c-Myc was targeted alone or in combination with other proteins, particularly BRAF, MEK and cyclin D1 (Figure 6A). We experimentally tested the key prediction from the network models. In order to target c-Myc expression, we treated melanoma cells with the BET bromodomain inhibitor, JQ1, as a single agent and in combination with MEKi (PD-0325901) or RAFi (vemurafenib). JQ1 directly targets bromodomains, especially those of the bromodomain protein BRD4, which marks select genes including *MYC* on mitotic chromatin. Inhibition of the BRD4 bromodomains with JQ1 downregulates *MYC* mRNA transcription and leads to G1 cell cycle arrest in diverse tumor types, such as multiple myeloma (Delmore et al., 2011; Loven et al., 2013; Puissant et al., 2013).

First, we asked whether we could affect c-Myc levels in SkMel-133 cells using JQ1. As measured by western blot experiments, c-Myc protein expression is reduced in response to JQ1 alone. c-Myc protein levels are further reduced when the cells are treated with a combination of JQ1 and MEKi or RAFi (Figure 6B).

To directly test the key prediction from the perturbation biology models, we measured the cell cycle progression response of melanoma cells to JQ1 in combination with the RAF and MEK inhibitors. We observed a strong synergistic interaction between JQ1 and RAFi (Figure 6C). 51% and 46% of melanoma cells are in G1 stage 24 hours after treatment with JQ1 (500nM) and RAFi (200nM) respectively, while 39% of cells are in G1 stage in the absence of any drug. On the other hand, when cells are treated with the combination of JQ1 and RAFi, a drastic increase in the fraction of cells arrested in G1-stage (84%) is observed. The single agent MEKi (50 nM) induces a strong G1-arrest phenotype in SkMel-133 cells (88% G1-stage in MEKi treated cells vs. 39% in non-drug treated cells.). Combination of MEKi with JQ1 arrests an even higher fraction of the cells (92%) in the G1-stage (Figure S5).

Before assessing the effect of JQ1-MEKi/RAFi combination on viability of melanoma cells (SkMel-133), we tested the effect of single agent JQ1 and found that the melanoma cells were considerably sensitive to single agent JQ1 treatment (cell viability IC50 = 200 nM). The sensitivity of SkMEl133 to JQ1 is similar to those of A375 and SkMel-5 lines (RAFi/MEKi sensitive, carrying *BRAFV600E* mutation) to another BRD4 inhibitor, MS417 (Segura et al, 2013). The observed sensitivity is also comparable to those of multiple myeloma and MYCN-amplified neuroblastoma cell lines, reported to be potentially JQ1-sensitive tumor types (Delmore et al., 2011; Puissant et al., 2013), and substantially higher than those of lung adenocarcinoma and MYCN-WT neuroblastoma cell lines (Lockwood et al., 2012; Puissant et al., 2013).

We tested the effect of combined targeting of c-Myc with MEK or BRAF on cell viability in SkMel-133 cells (Figure 6D). Strikingly, when combined with JQ1 (120nM), cell viability is reduced by 50% with 120 nM of RAFi (PLX4032), whereas the IC50 for single agent RAFi is >1 µM in RAFi-resistant SkMel-133 cells. Similarly, when combined with 5 nM MEKi (PD901), viability of SkMel-133 cells is reduced by 50% with 100 nM of JQ1, an IC50 value, which is close to those of the most sensitive multiple myeloma cell lines (Delmore et al, 2011). At higher doses (IC80), JQ1 is synergistic with both MEKi (Combination index, CI_85_ = 0.46) and RAFi (CI_85_ = 0.47) in SkMel-133 cells. At intermediate doses, JQ1 synergizes with RAFi (CI_50_ = 0.65) and has near additive interaction with the MEKi (CI_50_ = 0.85) (Figure 6E). Consistent with the observed synergy at high doses, both JQ1 combinations significantly improve the maximal effect level (A_max_, response to the drugs at highest doses), leading to lower cell viability beyond the levels reached by treatment with any of the agents alone. The observed improvement in A_max_ is particularly important since a subpopulation of cancer cells usually resist treatment even at highest possible drug doses. Treatments with drug combinations, such as those tested here will overcome or delay emergence of drug resistance if they can shrink the size of this resistant subpopulation (i.e. lead to improved A_max_).

## Discussion

We generated network models of signaling in melanoma cells to systematically predict cellular response to untested drug perturbations. Our modeling algorithm integrates high-throughput drug response profiles and pathway information from signaling databases. The scale and the predictive power of the models are beyond the reach of the currently available network modeling methods. Based on the predictions from models, we found that co-targeting MEK or BRAF with c-Myc leads to synergistic responses to overcome RAF inhibitor resistance in melanoma cells. Beyond nomination of effective drug combinations, the perturbation biology method paves the way for model-driven quantitative cell biology with diverse applications in many fields of biology.

### Cell type specificity

In network modeling, the experimental data provides the cell type-specific constraints while the priors introduce a probabilistic bias for background signaling information. Consequently, the network models are cell type specific and not only recapitulate known biology but also predict novel interactions. Moreover, the algorithm rejects a significant part of the interactions in the prior model. For example, the influences of cyclin E1 and cyclin D1 on RB phosphorylation are well known (Akiyama et al., 1992; Kato et al., 1993) and represented in the prior information model. The inferred models included the expected positive edge between cyclin D1 and RBpS807, but not between cyclin E1 and RBpS807. What are the genomic features in SkMel-133 cells that may lead to such context specific interactions? The gene product of *CDKN2A*, p16Ink4A, directly inhibits cyclin D1/CDK4 as it participates in a G1 arrest checkpoint (Serrano et al., 1993). On the other hand, the alternative *CDKN2A* gene product, p14ARF can inhibit the cyclin E1/CDK2 complex only indirectly through an MDM2/TP53/p21Cif1 dependent pathway and *CDKN2A* gene products appear to have no direct influence on cyclin E1 (Giono and Manfredi, 2007; Stott et al., 1998). In SkMel-133 cells, the homozygous deletion in *CDKN2A* most likely leads to excessive cyclin D1/CDK4 catalytic activity, which may override the influence of cyclin E1 /CDK2 complex on RBpS807. The descriptive aspect of the inferred models is of course inherently limited to the 99 components (nodes) included in models and constrained by data or prior information.

### Co-Targeting c-Myc and ERK signaling

Oncogenic alterations that decrease drug sensitivity may pre-exist in combinations or emerge sequentially in a tumor. In general, it is likely that tumors can escape therapy through alternative routes, rendering countermeasures difficult. One potential counter strategy is to identify and target proteins, on which multiple drug resistance pathways converge. Through quantitative simulations, we have predicted that c-Myc couples multiple signaling pathways such as MAPK and AKT to regulation of cell cycle arrest in SkMel-133 cells (Table 1). According to the predictions, co-targeting c-Myc with MEK, BRAF or cyclin D1 leads to the highest impairment in tumor growth. To test our predictions, we targeted c-Myc using the epigenetic drug JQ1, a BRD4 inhibitor that negatively effects c-Myc transcription. We treated cells with combinations of JQ1 and MEKi or RAFi. We showed that both combinations lead to synergistic cell viability response with particularly improved outcomes in high doses (lower Amax). A hallmark of high pharmacological efficacy is a drug’s ability to induce cellular response at doses sufficient to inhibit the immediate molecular targets. In SkMel-133 cells, however, the cell viability IC50s for RAFi and MEKi are at least one order of magnitude higher than the doses required to reduce the phosphorylation of immediate downstream targets (phospho-MEK and phospho-ERK, respectively) by 50% (Figure 6B, Table 2) and hence the cells are resistant to both drugs. When combined with JQ1, RAFi/MEKi doses, which are sufficient to reduce cell viability by 50%, are in the range of MEK and ERK phosphorylation IC50s (Table 2). Thus, the JQ1-MEKi/RAFi combinations shift the required doses to induce a cellular response close the doses required for inhibition of ERK pathway activity. Recently, it is reported that JQ1 and FLT3 tyrosine kinase inhibitor (TKI) combination shows synergy to overcome FLT3-TKI in AML cells (Fiskus et al., 2014). As far as we know, this is the first demonstration of a combined inhibition of a BET bromodomain protein and a protein kinase molecule to overcome drug resistance in solid tumors (Filippakopoulos and Knapp, 2014).

**Table 2.**
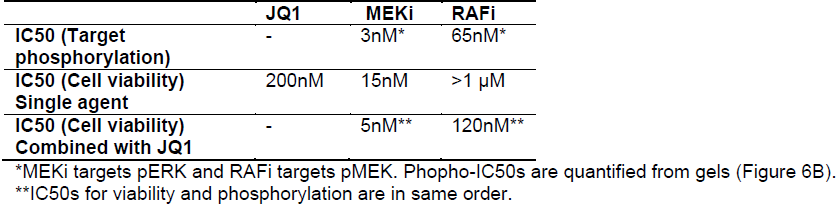
Drug resistance is overcome as the IC50 (cell viability) approaches the IC50 (target phosphorylation)

|  | JQ1 | MEKi | RAFi |
| --- | --- | --- | --- |
| IC50 (Target phosphorylation) | - | 3nM* | 65nM* |
| IC50 (Cell viability)<br>Single agent | 200nM | 15nM | >1 μM |
| IC50 (Cell viability)<br>Combined with JQ1 | - | 5nM** | 120nM** |

\*MEKi targets pERK and RAFi targets pMEK. Phospho-IC50s are quantified from gels (Figure 6B).
\*\*IC50s for viability and phosphorylation are in same order.

Targeting c-Myc can be highly toxic since c-Myc functions as a key transcription factor almost in all normal tissues and tumors (Soucek et al., 2008). MEK inhibitors are also highly toxic and their toxicity can only be managed with dose interruption and reduction (Flaherty et al., 2012). Use of synergistic drug combinations offers a potential solution to the drug toxicity problem as the drug doses in such combinations are substantially low compared to the doses in the single agent treatments. We conclude that co-targeting c-Myc and ERK pathway has the potential to go beyond single agent treatments in overcoming drug resistance and lowering drug toxicity in melanomas in genomic contexts similar to that in this study.

The model-based predictions provide comprehensive and testable hypotheses on complex regulatory mechanisms and development of novel therapies. The improvements in experimental data volume and signaling databases is likely to lead to network models with even higher predictive power. Coupling of the cell line-specific predictions to comprehensive genomic analyses will guide biologists to extrapolate the potential impact of the nominated combinations to tumors with similar genomic backgrounds. For example, we have developed genomics methods to classify tumors based on select oncogenic alterations and identify cell lines that most closely resemble tumors for further preclinical development (Ciriello et al., 2013; Domcke et al., 2013). By integrating network models, genomics and pathway analysis, one can expect to generate whole cell models of signaling and drug response in mammalian cells with potential applications to personalized medicine and genomically-informed clinical trials.

## Methods

### Cell cultures and perturbation experiments

The RAFi-resistant melanoma cell line SkMel-133 is used in all perturbation experiments. SkMel-133 cells are perturbed with 12 targeted drugs applied as single agents or in paired combinations (See Tables S1 and S3 for drug list, presumed targets, dosing and sources). In total, cells are treated with 89 unique perturbations. In paired combinations, each drug concentration is selected to inhibit the readout for the presumed target or the downstream effectors by 40% (IC40) as determined by Western blot experiments (Molinelli, Korkut, Wang et al., 2013) (Table S1). In single agent perturbations, each drug is applied at two different concentrations, IC40 and 2 x IC40. In validation experiments, (+)JQ1 (Cayman Chemicals) and the FDA approved RAFi PLX4032 (Selleckchem) are used.

### Reverse Phase Protein Arrays

Proteomic response profiles to perturbations are measured using reverse phase protein arrays (MD Anderson Cancer Center RPPA Core Facility). The cells are lysed 24 hours after drug treatment. Three biological replicates are spotted for each sample (i.e. drug condition) on RPPA slides. Each slide is interrogated with a particular antibody, so for each experimental condition 138 proteomic entities (levels of total protein or phoshoprotein) are profiled on 138 slides (Table S3).

### Quantitative phenotypic assays

All phenotypic measurements are made in perturbation conditions identical to those of proteomic measurements. Cell viability and cell cycle progression are measured using the Resazurin assay (72 hours after drug treatment) and flow cytometry analysis (24 hours after drug treatment) respectively. The percentage of cells in the G1, G2/M, and S phases and sub-G1 fraction are recorded based on the respective distribution of DNA content in each phase.

### Automated extraction of prior information from signaling databases

Pathway Extraction and Reduction Algorithm (PERA) was developed to automatically extract prior information from multiple signaling databases and generate a prior information network. The input to PERA is a list of (phospho) proteins identified by their HGNC symbols (*e.g. AKT1*), phosphorylation sites (*e.g.* pS473) and their molecular status (i.e., activating or inhibitory phosphorylation, total concentration). The output of PERA is a set of directed interactions between signaling molecules represented in a Simple Interaction Format (SIF). The PERA software is available at http://bit.ly/bp_prior as a free software under LGPL 3.0 (See supplementary methods for the details of the PERA).

### Mathematical description of network models

The network models represent the time behavior of the cellular system in a set of perturbation conditions as a series of coupled nonlinear ordinary differential equations (ODE) (Nelander et al., 2008).

**Equation 1. Network model ODEs**

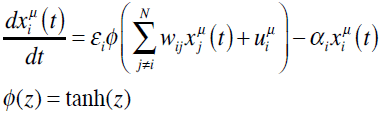

In the network models, each node represents the quantitative change of a biological variable, x_i^µ^_ [(phospho)protein level and phenotypic change] in the perturbed condition, µ, relative to the unperturbed condition. W_ij_ quantifies the edge strength, which is the impact of upstream node j on the time derivative of downstream node i. We assign semi-discreet values to each W_ij_, ***W***={*w_ij_,* ∀*w_ij_* ∈{-1,-0.8, …0.8,1}}. αi constant is the tendency of the system to return to the initial state, and εi constant defines the dynamic range of each variable i. The transfer function, *Φ*(x) ensures that each variable has a sigmoidal temporal behavior.

### Modified cost function for network inference with prior information

We have quantified the cost of a model solution by an objective cost function C(W). The network configurations with low cost represent the experimental data more accurately. Here, we have incorporated an additional prior information term to the cost function to construct models with improved predictive power. The newly introduced term in the cost function accounts for the prize introduced when the inferred wij is consistent with the prior information (See supplementary methods for the formulation of the modified cost with prior information term).

### Network model construction and response prediction

Network models are constructed with a two-step strategy. The method is based on first calculating probability distributions for each possible interaction at steady state with the Belief Propagation (BP) algorithm and then computing distinct solutions by sampling the probability distributions. We described the theoretical formulation, the underlying assumptions and simplification steps of the BP algorithm for inferring network models of signaling elsewhere (Molinelli, Korkut, Wang et al., 2013). The network models include 82 proteomic, 5 phenotypic and 12 activity nodes. Activity nodes couple the effect of drug perturbations to the overall network models (See Molinelli, Korkut, Wang et al. for quantification of activity nodes).

### Belief propagation

Belief propagation algorithm iteratively approximates the probability distributions of individual parameters. The iterative algorithm is initiated with a set of random probability distributions. In each iteration step, individual model parameters are updated (local updates) based on the approximate knowledge of other parameters, experimental constraints and prior information (global information). In the next iteration, the updated local information becomes part of the global information and another local update is executed on a different model parameter. The successive iterations continue over different individual parameters until the updated probability distributions converge to stable distributions. The iterations between the local updates and the global information create an optimization scheme that **W**= {w_ij_} is inferred given a probability model. Explicitly, the following cavity update equations are iteratively calculated until convergence.

**Equation 2. BP update equations**

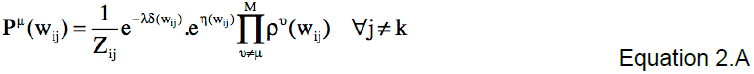

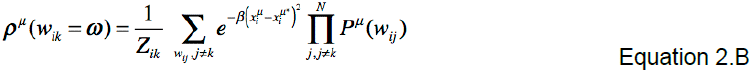

In Equation 2a, P^µ^(w_ij_) approximates the mean field of the parameters with a sparsity constraint (λδ(w_ij_)) and a bias from prior information restraints (η(wij)). In Equation 2b, ρ^µ^(w_ij_=ω) is a mean field derived change to the probability distribution of the locally optimized parameter, towards minimizing the model error (C^SSE^(W)). (See supplementary methods and Molinelli, Korkut, Wang et al. for derivation and implementation of BP equations).

### BP-guided decimation

Distinct networks models are instantiated from BP generated probability distributions with the BP-guided decimation algorithm (Figure S1) (Montanari et al., 2007). This procedure generates distinct and executable network models. In this study, 4000 distinct network models are generated in each computation.

### Simulations with in silico perturbations

Network models are executed with specific in silico perturbations until all system variables {xi} reach steady state. The perturbations acting on node i are exerted as real-valued uiµ vectors in model equation 1. The DLSODE integration method (ODEPACK) (Hindmarsh, 1993) is used in simulations (default settings with, MF = 10, ATOL= 1e-10, RTOL = 1e-20).

## Acknowledgements

We thank N. P. Gauthier, P. Kaushik, M.L. Miller, D.S. Marks, A. Braunstein, A. Pagnani, R. Zecchina, M. Cokol, and N. Rosen for helpful discussions. This work was funded in part by the Center for Cancer Systems Biology grant U54 CA148967 (NIH), the National Resource for Network Biology grant GM103504 (NIH), the Research Resource for Biological Pathways grant U41 HG006623 (NHGRI) and a Melanoma Research Alliance Established Investigator Award.

